# Cytotoxic effects of tRNA-derived fragments and tRNA halves from non-pathogenic *Escherichia coli* strain on colorectal cancer cells

**DOI:** 10.1101/2020.05.18.103366

**Authors:** Kai-Yue Cao, Yu Pan, Tong-Meng Yan, Peng Tao, Yi Xiao, Zhi-Hong Jiang

**Author notes:** Corresponding author: Dr. Zhi-Hong Jiang, State Key Laboratory of Quality Research in Chinese Medicine, Macau University of Science and Technology, Avenida Wai Long, Taipa, Macau SAR, China. (Zhi-Hong Jiang).

## Abstract

Transfer RNAs (tRNAs) purified from non-pathogenic *E. coli* strain (NPECS) possess cytotoxic properties on colorectal cancer cells. In the present study, the bioactivity of tRNA halves and tRNA fragments (tRFs) derived from NPECS are investigated for their anticancer potential. Both tRNA halves and tRF mimics studied exhibited significant cytotoxicity on colorectal cancer cells, with the latter being more effective suggesting that tRFs may be important contributors to the bioactivities of tRNAs derived from gut microbiota. Through high-throughput screening, EC83 mimic, a double-strand RNA with a 22 nt 5’-tRF derived from tRNA-Leu(CAA) as antisense chain, was identified as one with the highest potency (IC_50_=52 nM). Structure-activity investigations revealed that 2’-*O*-methylation of the ribose of guanosine may enhance the cytotoxic effects of EC83 mimic *via* increasing the stability of its tertiary structure. Consistently, 4-thiouridine substitution reversed this increased stability and the enhanced cytotoxic effects. This provides the first evidence that the bioactivity of tRF mimics would be impacted by chemical modifications. Furthermore, the present study provides the first evidence to suggest that novel tRNA fragments derived from the gut microbiota may possess anti-cancer properties and have the potential to be potent and selective therapeutic molecules.

**IMPORTANCE:** While the gut microbiota has been increasingly recognized to be of vital importance to human health and disease, the current literature shows that there is a lack of attention given on the non-pathogenic *Escherichia coli* strain. Moreover, the biological activities of tRNA fragments (tRFs) derived from bacteria have rarely been investigated. The findings from this study revealed tRFs as a new class of bioactive constituents derived from gut microorganisms, suggesting that studies on biological functional molecules in intestinal microbiota should not neglect tRFs. The research of tRFs would play an important role in biological research of gut microorganisms, including bacteria-bacteria interaction, gut-brain axis, gut-liver axis, etc. Furthermore, the guidance on the rational design of tRF therapeutics provided in this study indicates that further investigations should pay more attention to these therapeutics from probiotics. The innovative drug research of tRFs as potent druggable RNA molecules derived from intestinal microorganisms would open a new area in biomedical sciences.

## Introduction

The microbiome in the human gastrointestinal tract consisting trillions of microorganisms comes into being within days after birth (1). Recent studies revealed that the gut microbiota in general provides beneficial effects to the host by aiding gastrointestinal and immune functions (2). While it can promote health, it may also sometimes cause diseases. For examples, non-alcoholic fatty liver, type 2 diabetes, inflammatory bowel disease, obesity, and colorectal cancer have been linked to the microbiota (3). *Escherichia coli* (*E. coli*) is a gram-negative gut commensal bacteria found in the colon of mammals and reptiles (4, 5, 6). It has become one of the most important model organisms due to its fast growth in chemically defined media *in vitro. E. coli* can be categorized into four main phylogenetic groups, A, B1, B2 and D. Pathogenic *E. coli* strains (PECS) from group B2 have been found to cause gastroenteritis, neonatal meningitis, hemorrhagic colitis, urinary tract infections, Crohn’s disease, etc. Advanced studies have revealed that PECS containing peptide synthetase-polyketide synthetase (pks) island could induce DNA double strand breaks in human cell lines, and thus PECS may be able to accelerate the development of colorectal cancer (7-9), which is a leading cause of cancer-related deaths worldwide (10, 11). On the other hand, *E. coli* strains of groups A and D are not known to cause diseases and can even be beneficial to their hosts, e.g. by producing vitamin K_2_ (5). Thus, the functions of these non-pathogenic *E. coli* strains (NPECS) demand attention from researchers.

In one of our previous studies, tRNA-Val(UAC) and tRNA-Leu(CAG), purified from NPECS by two-dimensional liquid chromatography, have been shown to exhibit significant cytotoxicity on HCT-8 human colorectal cancer cells (12). As the most abundant class of small RNAs (<200 nucleotides) (13), tRNAs have been revealed to regulate RNA splicing, RNA translation and DNA replication (14). It has been reported that endogenous tRNAs may be cleaved by multiple ribonucleases, and under stress conditions, may produce tRNA-derived fragments (tRFs) and tRNA halves (15-18) which have been associated with broad biological functions, such as miRNA-like regulation of protein translation and cellular stress responses (19, 20). In addition, endogenous tRFs can suppress human breast cancer through targeting YBX1, suggesting possible application as pharmacological agents (21). On the other hand, tRNA halves were found to suppress breast cancer *via* targeting FZD3 (22). Therefore, it is reasonable to hypothesize that the cytotoxic effects of tRNA-Val(UAC) and tRNA-Leu(CAG) from NPECS reported in our previous paper (12) may be mediated by the formation of tRFs and tRNA halves. Therefore, we embarked on a study to evaluate and compare tRNA halves and tRF mimics (double-strand RNA with tRF as antisense chain) derived from these two RNAs in terms of cytotoxicity on colorectal cancer cells. Moreover, the structure-cytotoxicity activity relationship of tRF mimics was studied using chemically modified tRF mimics.

## Results

### Cytotoxicity evaluations

tRNA halves of tRNA-Val(UAC) and tRNA-Leu(CAG) were prepared from intact tRNAs (Fig. S1) by S1 nuclease, which cleaves the tRNAs into two half fragments at their anticodon loop (Fig. 1A-C). Cytotoxicity studies, using HCT-8 cells with liposomal transfection at a concentration of 50 nM, showed that the intact tRNA-Val(UAC) and tRNA-Leu(CAG) and their tRNA halves and tRF mimics (double-strand RNAs with 22 nt-long 5’- and 3’-tRF as antisense chain) significantly decreased cell viability of HCT-8 cells by various extents ranging from approximately 36-80%, with the exception of 5’-tRNA half of tRNA-Val(UAC) (Fig. 1D, E). However, the 5’-tRF mimics of both tRNAs are the most effective RNA.

**Figure 1.**
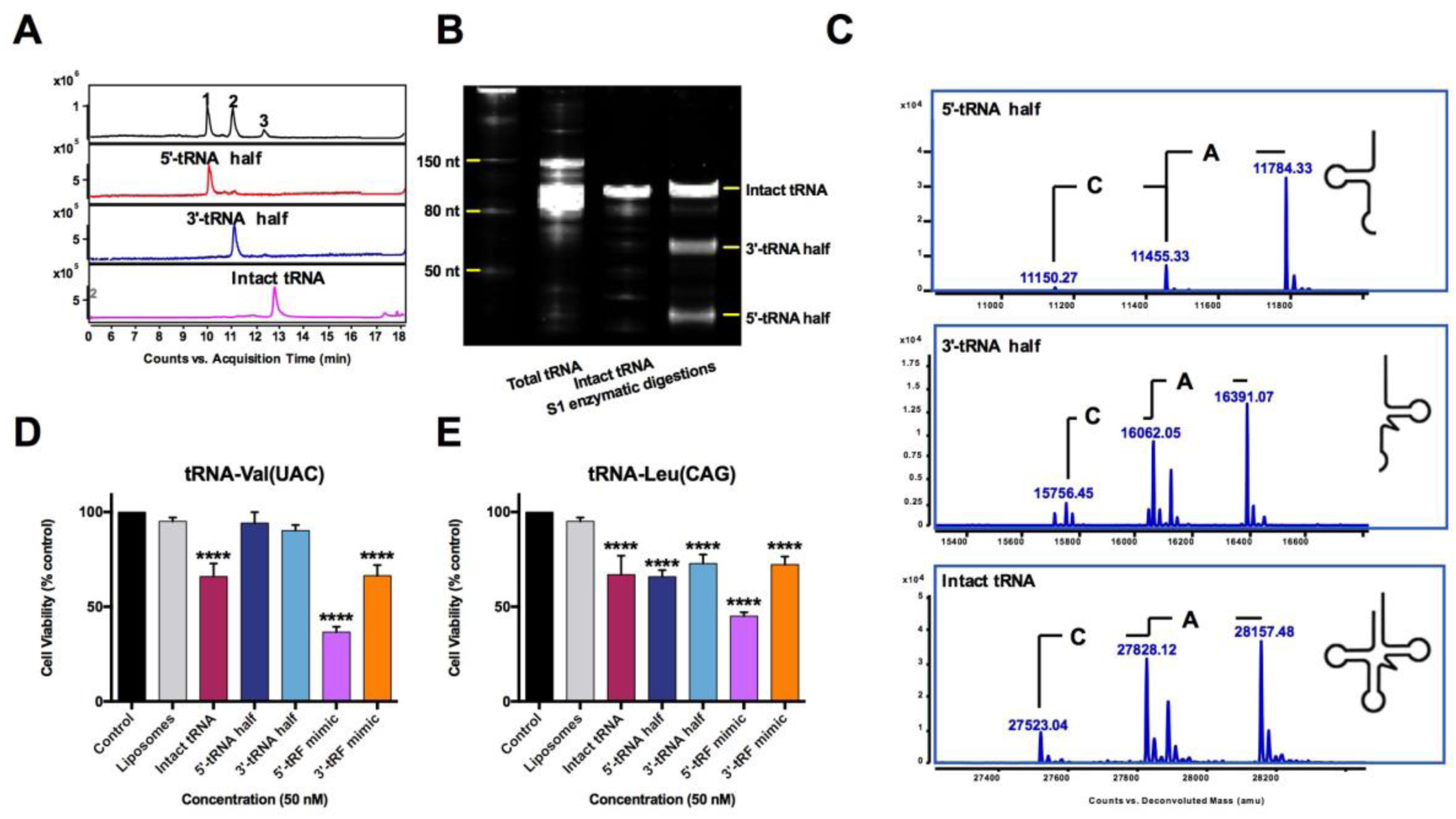
The cytotoxicity of tRF mimics is significantly stronger than that of tRNA halves on HCT-8 cells. (**A**) Typical UHPLC chromatogram of 5’-and 3’-tRNA halves of tRNA-Leu(CAG) under UV 260 nm. (**B**) Urea-polyacrylamide gel electrophoresis of 5’-and 3’-tRNA halves of tRNA-Leu(CAG), confirming that the purified tRNA was successfully digested into two products. (**C**) UHPLC-MS analysis of 5’-and 3’-tRNA halves of tRNA-Leu(CAG) showing that the molecular weights of digestion products were in accordance with 5’- and 3’-tRNA halves of *E. coli* tRNA. (**D**) Cytotoxic comparison of tRNA halves, tRF mimics and individual tRNAs of tRNA-Val(UAC) on HCT-8 cells. (**E**) Cytotoxic comparison of tRNA halves, tRF mimics and individual tRNAs of tRNA-Leu(CAG) on HCT-8 cells. Data are shown as the means ± SD of three independent experiments. ****, *P*<0.0001 (one-way ANOVA followed by post hoc analysis).

### High-throughput screening of *E. coli* tRF mimics

A total number of 82 tRFs, including 5’-tRF and 3’-tRF with 22 nt, the most abundant types of tRFs (23), were selected as antisense strands of tRF mimics in siRNA form, and their cytotoxic effects were screened using HCT-8 cells (Fig. 2A). Following an initial screen at 50 nM, a total of 12 tRF mimics which exhibited the highest effects were selected for detailed analysis. Fig. 2B shows the dose-dependent cytotoxic effects of these 12 tRF mimics. The most effective tRF mimics is EC83 mimic [a double-strand RNA with a 22 nt 5’-tRF derived from tRNA-Leu(CAA) as antisense chain] with an IC_50_ value of 52.8 nM (Fig. 2C).

**Figure 2.**
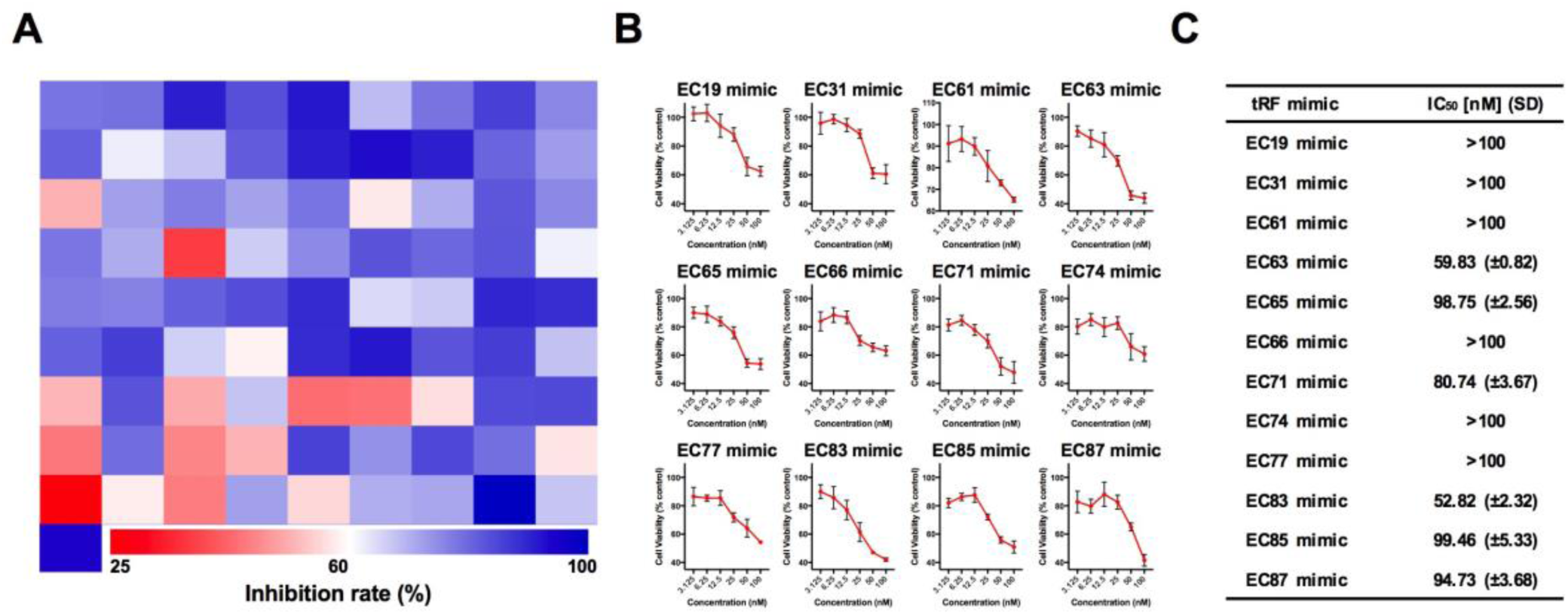
High-throughput screening hit EC83 mimic with the strongest cytotoxicity than other NPECS tRF mimics on HCT-8 cells. (**A**) Heatmap of high-throughput screening of 82 tRF mimics hit 12 tRF mimics with the potency of reducing cell viability of HCT-8 cells. (**B**) Dose-dependent investigations of top 12 hit tRF mimics. (**C**) IC_50_ values with standard deviations of top 12 hit tRF mimics, showing EC83 mimic has the strongest cytotoxicity against HCT-8 cells.

### Structure-activity relationship of tRF mimics

It has been demonstrated that chemical modifications of tRNA can alter the biological functions of tRFs (24). According to the reported modifications of 2’-*O*-methylguanosine (Gm) and 4-thiourdine (s^4^U) in tRNA-Leu(CAA), 3 modified derivatives of EC83 were designed and synthesized, namely EC83-M1 (s^4^U), EC83-M2 (Gm) and EC83-M3 (s^4^U+Gm) mimics. Fig. 3A shows the LC-MS analysis which confirmed good agreement of their molecular weights to the theoretical values (Fig. 3B) (25). Fig. 4A shows that EC83 mimic and its derivatives decreased cell viability of colorectal cancer cells (HCT-8 and its 5-FU-resistant strain, LoVo and its 5-FU-resistant strain). Notably, EC83-M2 mimic exhibited the strongest cytotoxicity with lowest IC_50_ value among four tRF mimics in the four cancer cell lines (Fig. 4B). These results indicated that 2’-*O*-methylation of the ribose of guanosine may increase cytotoxicity of the EC83 mimic, which may be reversed by the 4-thiouridine modification. In comparison, 5-FU has IC_50_ values in the μM concentration range, which is more than 700 times of those of EC83-M2 mimic, indicating this tRF mimic’s extraordinary cytotoxicity toward cancer cells. It is noteworthy that all tRF mimics showed the same potency in the 5-FU resistant cells as in the non-resistant cells.

**Figure 3.**
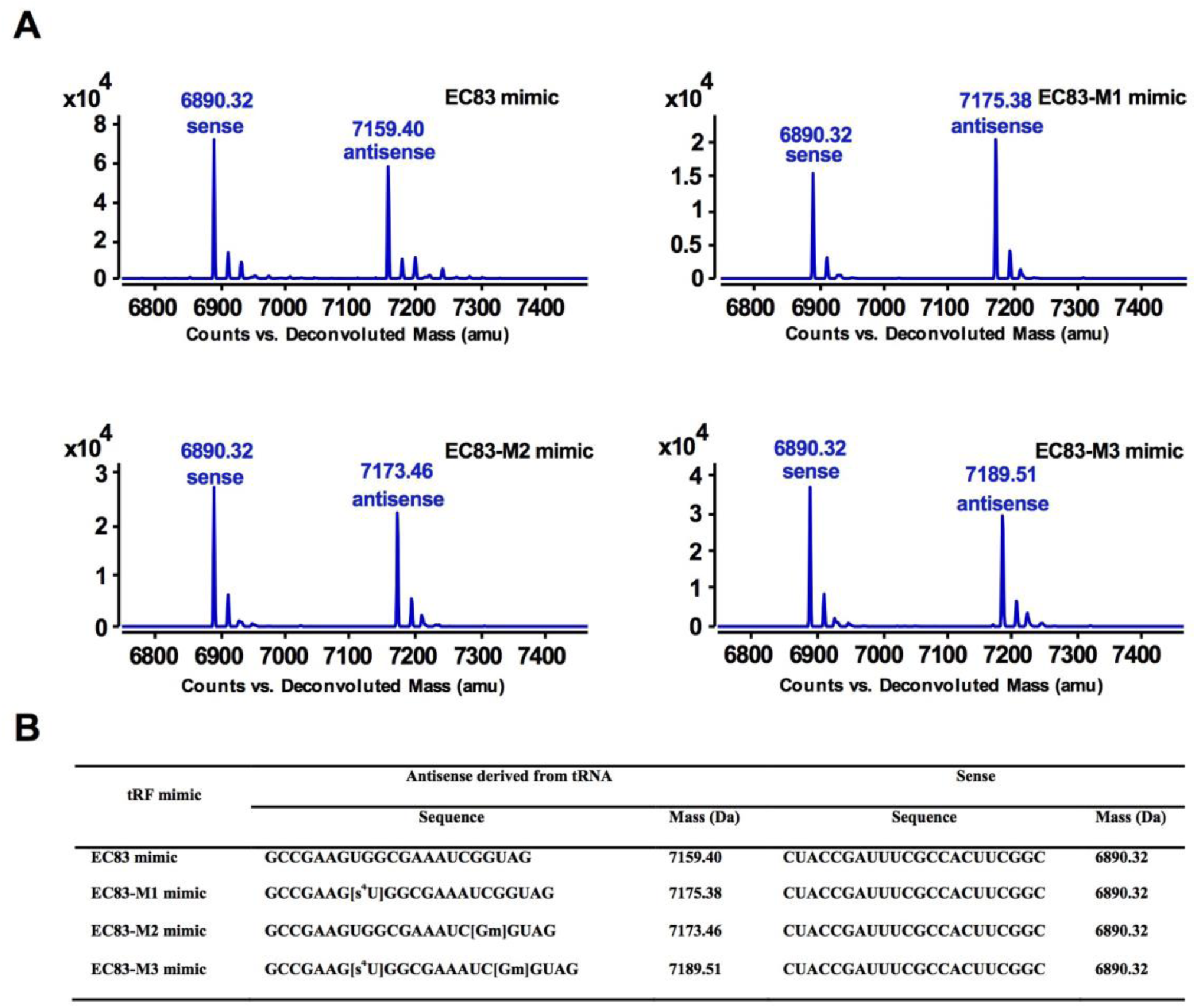
UHPLC-MS analysis confirmed the accurate molecular weights of EC83 mimic and its chemically modified derivatives. (**A**) UHPLC-MS analysis of EC83 mimic and its modified derivatives confirmed the agreement of their molecular weights to the theoretical values. (**B**) Sequence information and deconvolution MS of EC83 mimic and its modified derivatives.

**Figure 4.**
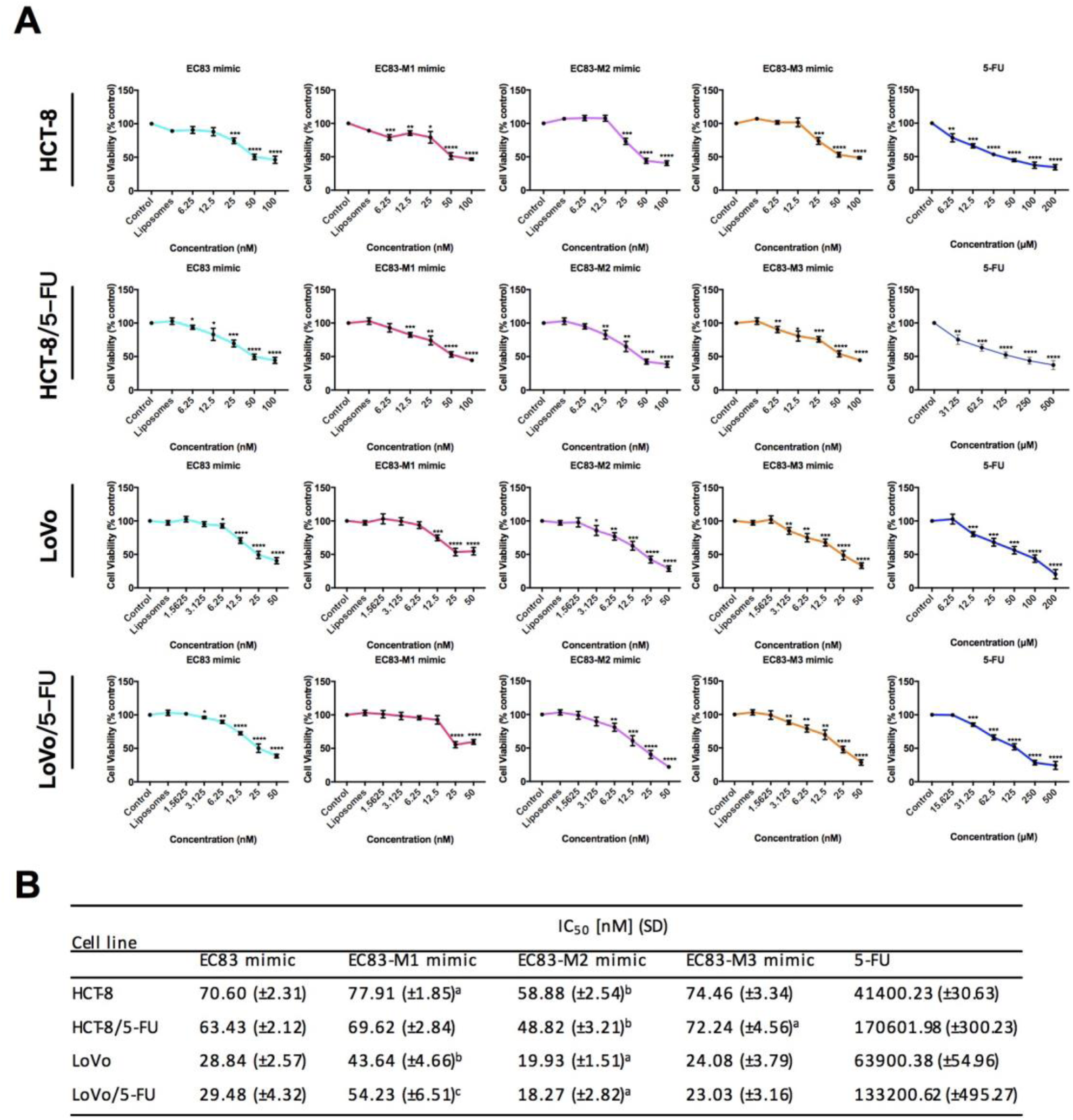
Gm may increase the cytotoxicity of EC83 mimic on colorectal cancer cells and their 5-FU resistant cells, which may be reversed by s^4^U. (**A**) Dose-dependent investigation of EC83 mimics and 5-FU on HCT-8, 5-FU resistant HCT-8, LoVo and 5-FU resistant LoVo cells. *, *P*<0.05; **, *P*<0.01; ***, *P*<0.001; ****, *P*<0.0001 (two-tailed Student t test). (**B**) IC_50_ values with SD of EC83 mimics and 5-FU, demonstrating EC83-M2 mimic exhibit stronger cytotoxicity than EC83 mimic, while EC83-M1 mimic has a worse cytotoxicity. a, *P*<0.1; b, *P*<0.05; c, *P*<0.01 (one-way ANOVA followed by post hoc analysis).

### Clonogenic assay

The effectiveness of EC83 mimic and its modified derivatives in preventing colony formation in the clonogenic assay was performed using HCT-8 (EC83 mimics at 50 nM and 5-FU at 50 μM) and LoVo cells (EC83 mimics at 25 nM and 5-FU at 50 μM) (Fig. 5A, B). The results indicated that all tRF mimics together with 5-FU are very effective in suppressing clonogenic ability in both cell lines with colonies reduced to about 1-19 % of control. Compared to EC83 mimic treated HCT-8 cells, EC83-M2 mimic treated ones exhibited a lower clonogenic ability, while EC83-M1 mimic treated ones have a higher survival percentage. On LoVo cells, EC83-M2 mimic treated cells exhibited a comparable clonogenic ability to those treated by EC83 mimic, while EC83-M1 mimic treated ones have a significantly higher survival percentage. It is noteworthy that EC83-M3 mimic treated cells (both HCT-8 and LoVo) have lower survival percentages than those of EC83-M1 mimic treated cells and higher survival percentages than those of EC83-M2 mimic treated cells. As a positive control, 5-FU exhibited a low clonogenic ability (10.6±5.0% on HCT-8 cells and 8.6±6.4% on LoVo cells). Overall, these results were in accordance with those of MTT assay, suggesting that 2’-*O*-methylguanosine might enhance the cytotoxic effectiveness of EC83 mimic, while 4-thiouridine has an opposite effect.

**Figure 5.**
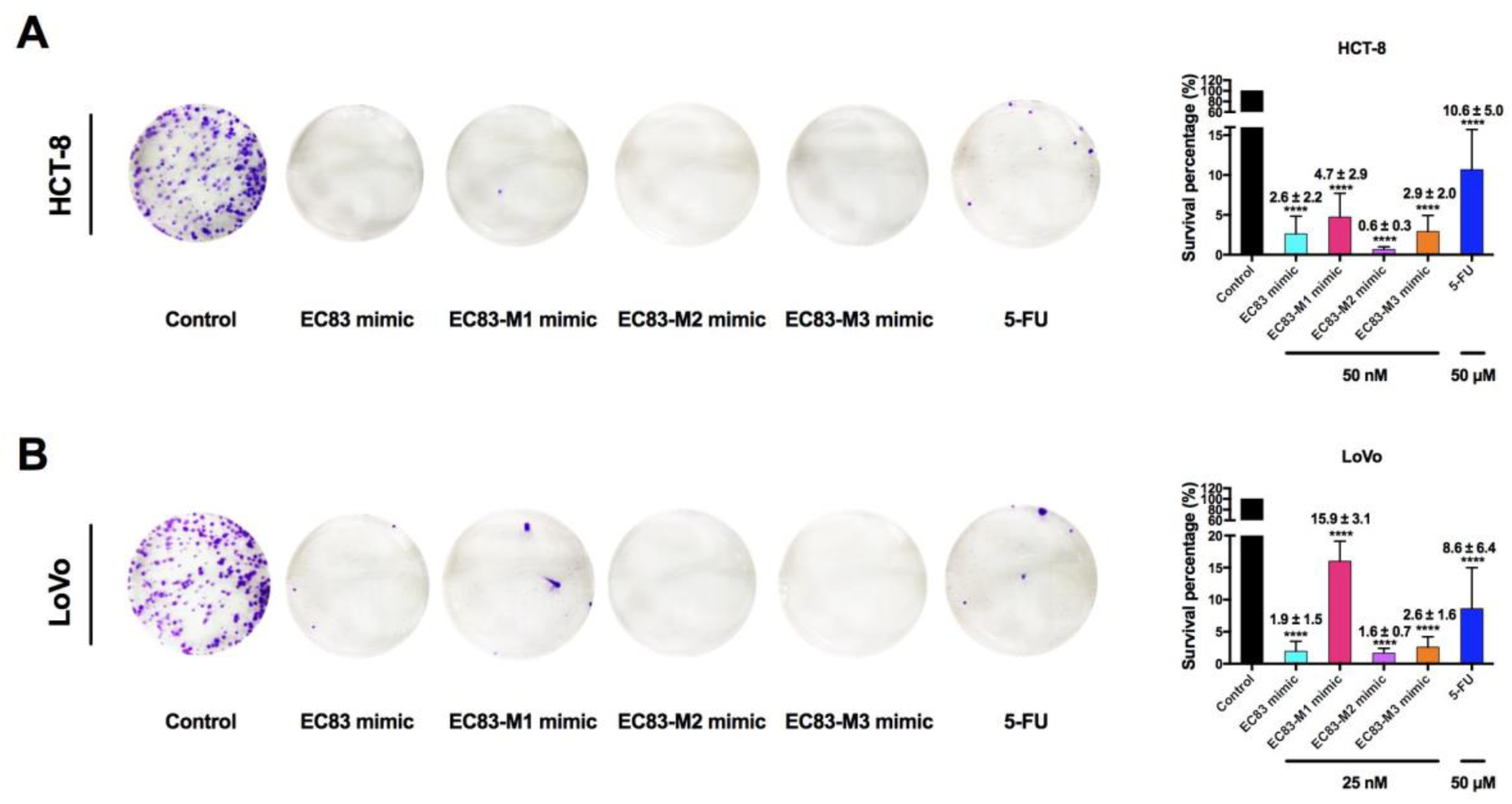
Gm might enhance the cytotoxic effectiveness of EC83 mimic, while s^4^U might has an opposite effect. (**A**) Clonogenic assay of EC83 mimic, EC83-M1 mimic, EC83-M2 mimic and EC83-M3 mimic on HCT-8 and LoVo cells. (**B**) Survival percentages of HCT-8 and LoVo cells treated by EC83 mimic and its modified derivatives. Data are shown as the means ± SD of three independent experiments. ****, *P*<0.0001 (two-tailed Student t test).

### Wound healing assay

The migration of cancer cells is important to cancer development. As shown in Fig. 6, at 24 and 48 h, EC83-M2 mimic treated HCT-8 cells at 50 nM (−38.6±10.6% and −48.0±5.3%) exhibited significant less wound healing rate (WHR) than EC83 mimic (4.0±2.4% and 2.8±5.3%), while EC83-M1 mimic treated ones have a higher WHR at the same concentration (19.4±2.1% and 11.4±2.8%). On LoVo cells, EC83-M2 mimic treated cells at 25 nM (−4.3±4.0% and −1.9±2.7%) exhibited a comparable WHR to EC83 mimic (−2.6±5.1% and −1.6±2.1%) at 24 and 48 h, while EC83-M1 mimic treated ones have a significant higher WHR at the same concentration (6.8±3.7% and 16.3±2.9%). Meanwhile, EC83-M3 mimic treated cells have less WHR than those of EC83-M2 mimic treated cells. As a positive control, 5-FU treated HCT-8 cells (11.7±4.2% and 13.8±2.9%) and LoVo cells (10.1±2.6% and −9.5±3.0%) at 50 μM exhibited a low HR on at 24 and 48 h. These results demonstrated that 2’-*O*-methylation of the ribose of guanosine may increase the inhibition effect on ability of wound healing of colorectal cancer cells of EC83 mimic, while 4-thiouridine might have an opposite effect, which were in accordance with the above results.

**Figure 6.**
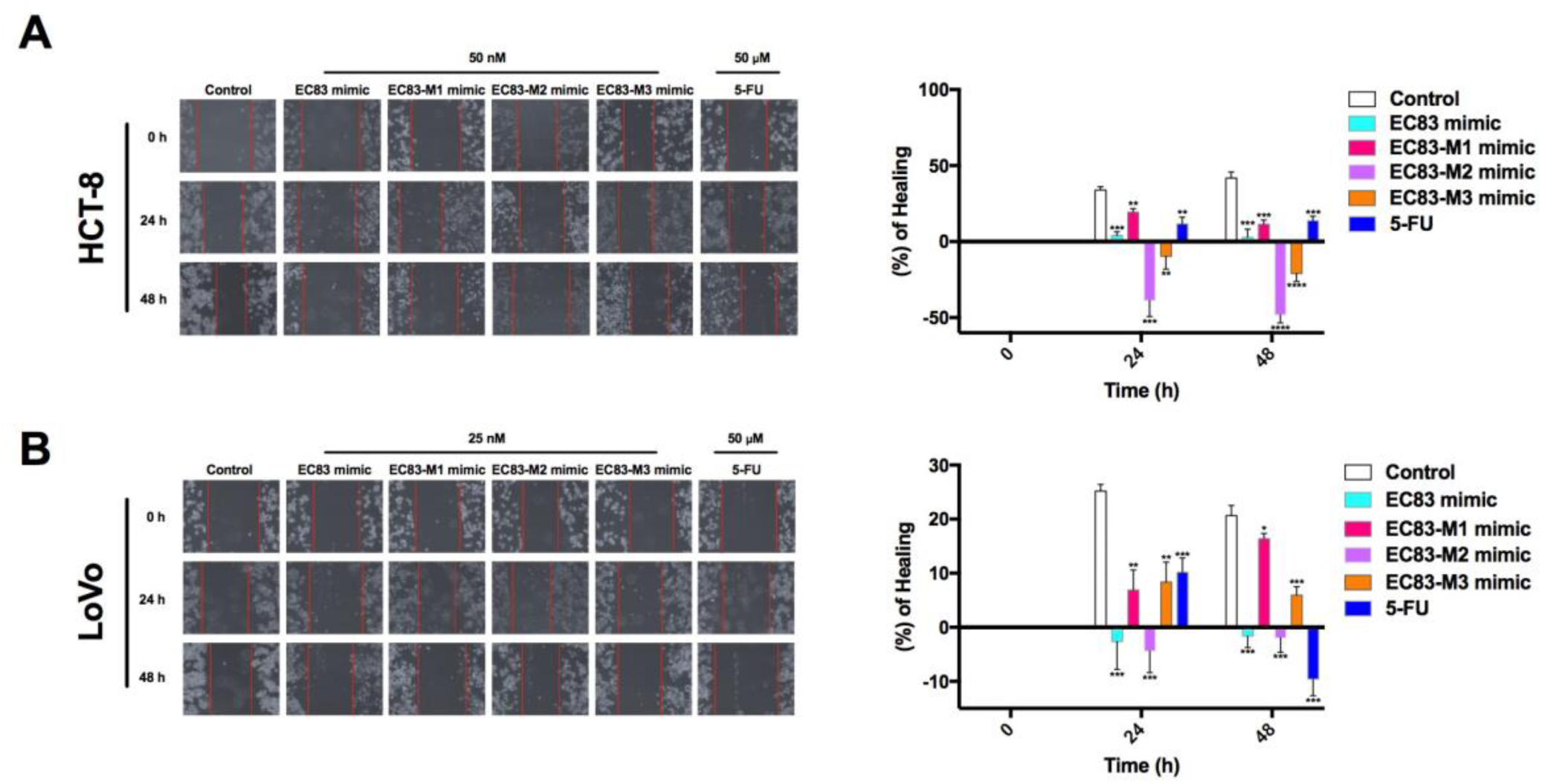
Gm may increase the inhibition effect of EC83 mimic on ability of wound healing of colorectal cancer cells, while s^4^U might has an opposite effect. (**A**) Wound healing assay of EC83 mimic, EC83-M1 mimic, EC83-M2 mimic and EC83-M3 mimic on HCT-8 and LoVo cells. (**B**) Wound healing rate of HCT-8 and LoVo cells treated by EC83 mimic and its modified derivatives. Data are shown as the means ± SD of three independent experiments. *, *P*<0.05; **, *P*<0.01; ***, *P*<0.001; ****, *P*<0.0001 (two-tailed Student t test).

### Evaluation of structural stability of EC83 mimics

To evaluate the stability of EC83 mimics at the molecular level, their structures were initially assumed to be the standard A-form double helix for molecular dynamics (MD) simulations. Starting from each of these initial structures, 100 ns simulation at 300 K in NPT ensemble revealed that EC83 mimic remained as A-form double helix, while the three modified derivatives transformed into more compact helical conformations rapidly (Fig. 7). Moreover, the fluctuation of the root-mean-square deviation (RMSD) of EC83 mimic is significantly larger than the others, indicating that their transformed helical structures are more stable than the A-form standard helix (Fig. 8).

**Figure 7.**
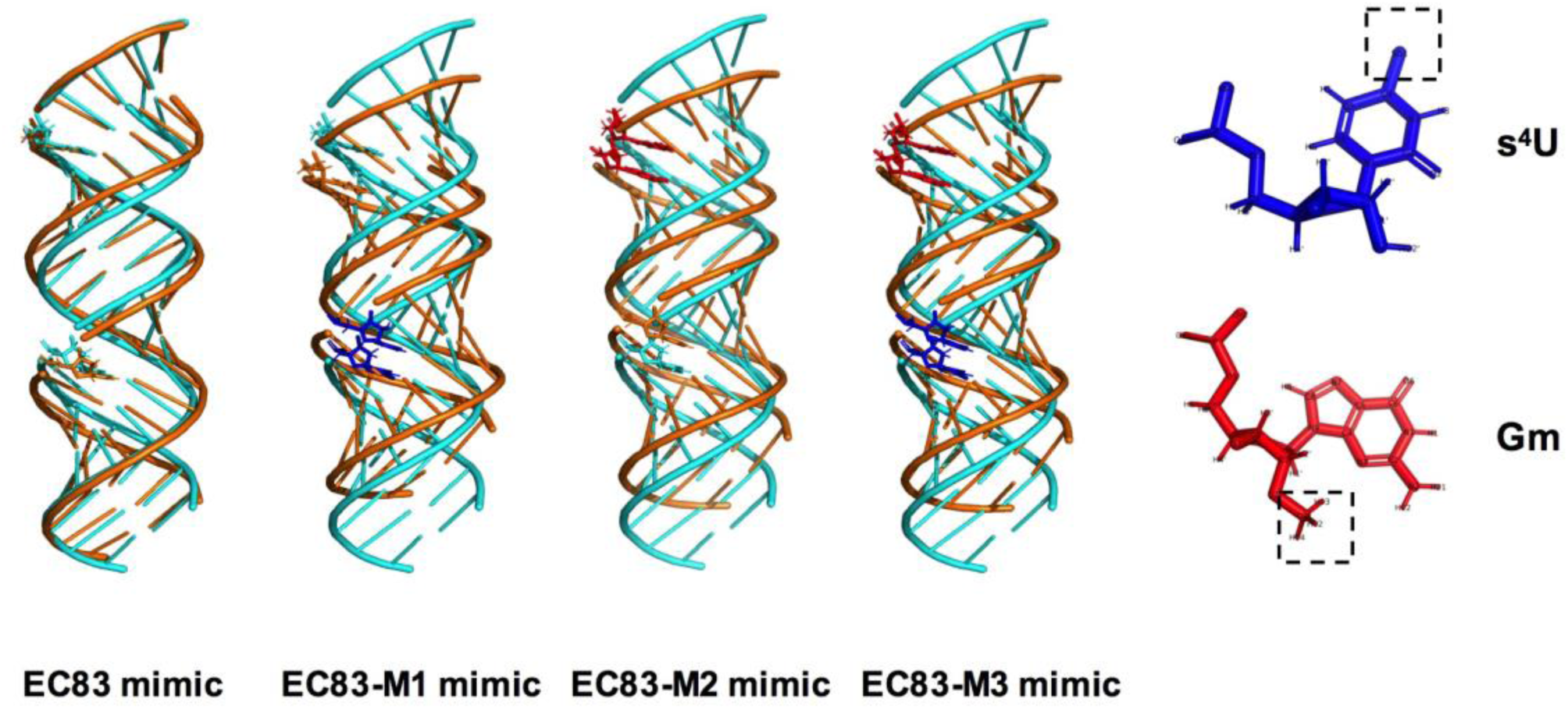
Simulated 3D initial structure (cyan) and transformed structure (gold) of EC83 mimic and its chemically modified derivatives.

**Figure 8.**
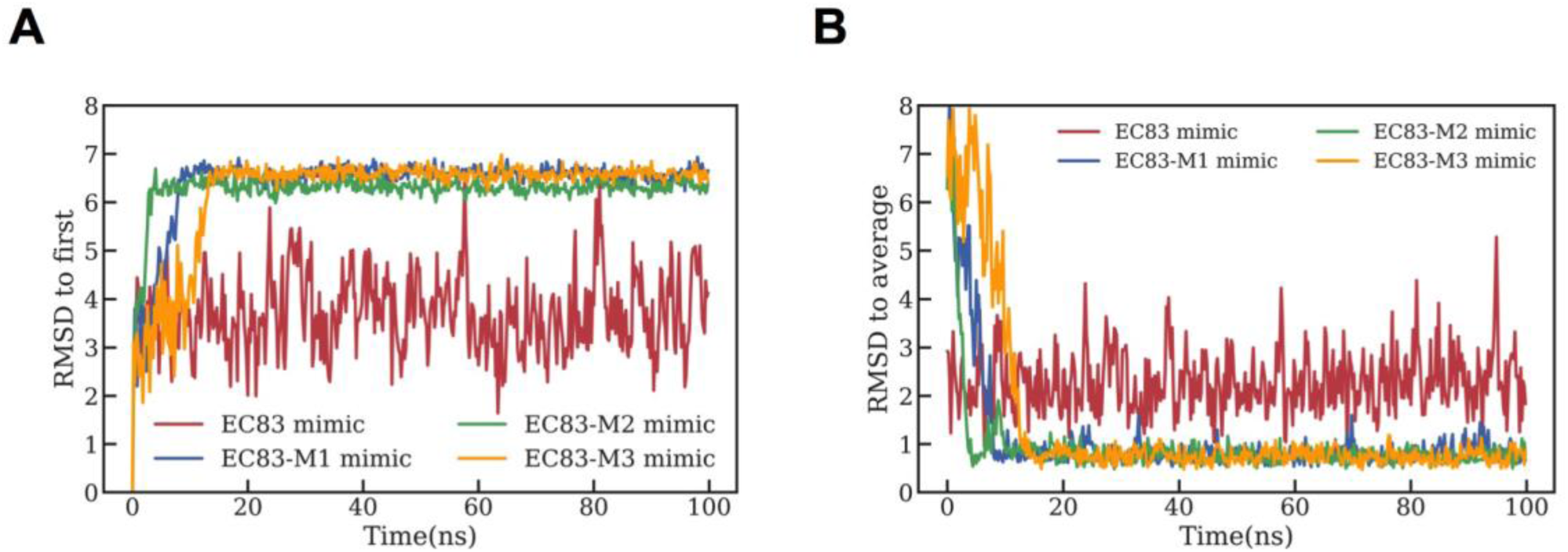
Chemically modified EC83 mimics are structurally more stable than unmodified EC83 mimic. (**A**) RMSD to the initial A-form structure and (**B**) stable structure for four dsRNAs changing over time.

Further analysis by 200 ns simulations with temperature increasing gradually from 300 K to 500 K (interval=10 K), the mimics, as shown in Fig. 9A, lost their stable structures in the order of EC83 mimic, EC83-M1 mimic or EC83-M3 mimic, and EC83-M2 mimic. Here, the unfolding structure was defined by their RMSD value above 5 Å relative to the stable structure. Further shown in Fig. 9B, the hydrogen bond (Hbond) ratio of EC83-M2 mimic remained above 40% at 260 ns, while the other 3 were under 20%. These results demonstrated that EC83-M2 mimic is structurally the most stable of the derivatives which are also more stable than the unmodified EC83 mimic, suggesting that 2’-*O*-methylation of guanosine enhances the tertiary structural stability of EC83 mimic. On the other hand, s^4^U appears to reverse the stabilizing effect of Gm. The results suggest that the derivatives are structurally more stable than the EC83 mimic, and that EC83-M2 is the most stable among the three derivatives.

**Figure 9.**
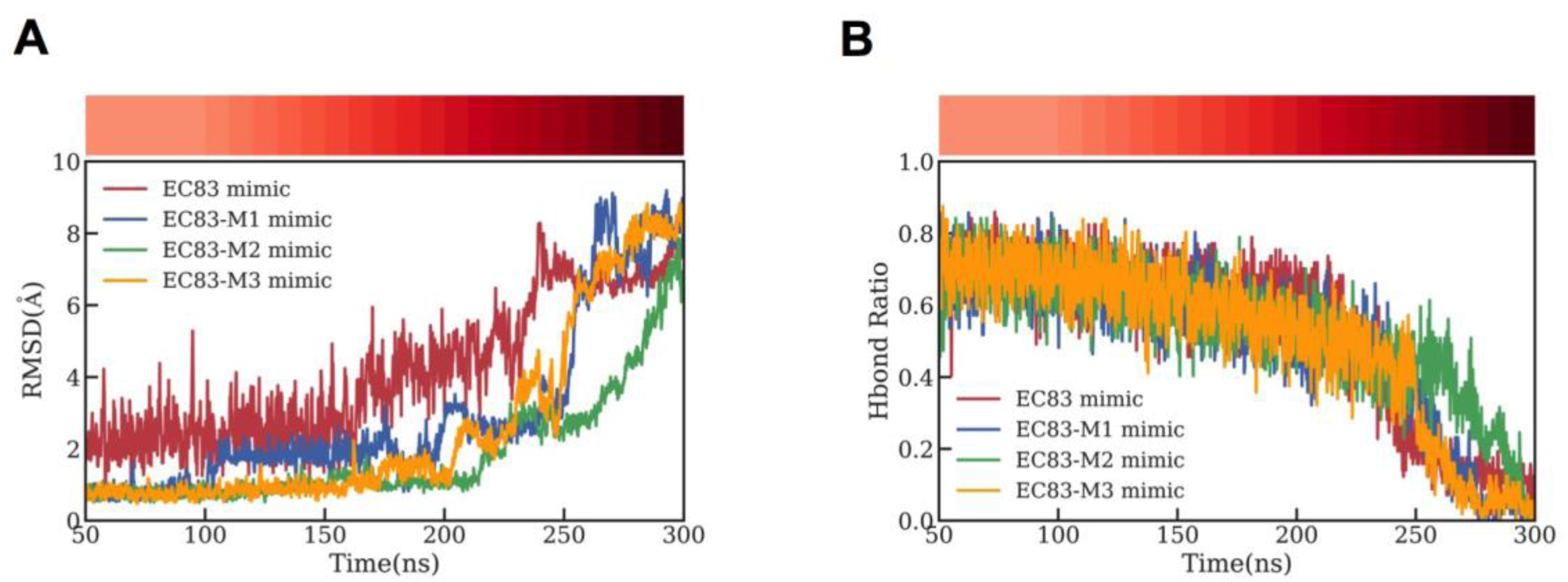
Gm enhances the tertiary structural stability of EC83 mimic, while s^4^U appears to reverse the stabilizing effect of Gm. With the increasing of temperature, the RMSD (**A**), using the stable structure as references, and the hydrogen bond (Hbond) ratio (**B**) of four dsRNAs changing over time. The temperature at each time point is shown in the colorbar above.

## DISCUSSION

Colorectal cancer (CRC) has become the second most frequently diagnosed malignancy worldwide accounting for ∼500,000 deaths each year. Risk factors of CRC include family history, diet high in red and processed meat, heavy alcohol intake, smoking, overweight or obesity, age (being older) and inflammatory bowel disease (26). Currently, cytotoxic chemotherapy is widely used as monotherapy or a component of combined therapy for CRC treatment (27). However, development of resistance to treatment is a big challenge. RNAs are able to regulate gene expression and therefore may have the potential to be a new class of cancer therapeutics which are non-toxic and have desirable selectivity for a specific target gene (28). The present study describes, for the first time, the ability of tRNA halves and tRFs derived from NPECS that exhibit cytotoxic activity against colorectal cancer cells. They are very effective with IC_50_ at the 10^−8^M concentration range, as compared to the 10^−4^M range of 5-FU. Interestingly, tRFs exhibited higher potency than tRNA halves, suggesting that tRFs might play an important role in the cytotoxic effects of tRNAs derived from gut microbiota. Importantly, the more potent EC83 mimic and its modified derivatives were observed to be just as effective on 5-FU-resistant CRC cells as on non-resistant cells, indicating that microbiota-derived tRFs may be an effective treatment even in CRC resistant to conventional chemotherapy. This observation supports the idea that the cytotoxic actions of tRFs differ from that of 5-FU. The exact mechanism of action remains to be investigated.

As a class of highly modified nucleic acid (29-34), tRFs have been proven to possess miRNA-like regulatory function in relation to RNA interference (24). Functional improvements have been reported following chemical modifications of tRFs (35). The loss of cytosine-5 RNA methylation was found to enhance the angiogenin-mediated endonucleolytic cleavage of tRNA leading to an accumulation of 5’-tRFs, and subsequently to reduced cell size and increased apoptosis (36). Consistently, the present results revealed that 2’-*O*-methylation of guanosine enhanced the cytotoxic effects of EC83 mimic perhaps *via* the observed increased stability of the 3D-structure. This was supported by the observation that 4-thiouridine substitution reversed this increased stability as well as the enhanced cytotoxic effects. This first structure-activity investigation provided evidence that the bioactivity of tRF mimics can be impacted by chemical modifications. Further structural analysis is required to decipher the critical structural differences between these modified tRF mimics.

Recently, it has been reported that colonic bacteria, especially PECS, bear a close relationship with CRC development (37, 38). Thus, current interests in clinical studies focus on PECS, such as EHEC O157:H7 which produces metabolites or toxins that can cause DNA damage in colon and CRC development (39-41). As the genome size and gene number of NPECS (3.98 Mb, 3696) differ from that of PECS (5.86 Mb, 5919) (42), NPECS and PECS may produce completely different profiles of tRNA fragments through different chemical modifications. This may raise the possibility of identifying CRC biomarker from tRNA molecules derived from PECS.

In summary, these findings revealed tRNA halves and especially tRFs as new class of bioactive constituents derived from gut microorganisms themselves and put new insights into their therapeutic effects on human diseases, which broadened our knowledge of beneficial effects of gut microbiota. Also, the present study demonstrates that studies on biological functional molecules in intestinal microbiota should not neglect tRFs since they are bioactive constituents (21). The research of tRFs would play an important role in biological research of gut microorganisms, including bacteria-bacteria interaction, gut-brain axis, gut-liver axis, etc. Furthermore, with the increasing interest in the identification of tRFs in bacteria (43, 44), the guidance on the rational design of tRF therapeutics provided in this study suggests that further investigations should pay more attention to these therapeutics from probiotics. The innovative drug research of tRFs as potent druggable RNA molecules derived from intestinal microorganisms would open a new area in biomedical sciences.

## MATERIALS AND METHODS

### Chemicals and reagents

*Escherichia coli* MRE600 total transfer ribonucleic acid was purchased from Roche (Switzerland). Biotin labeled single-stranded DNA oligonucleotides were obtained from B.G.I., China. Low range ssRNA ladder was purchased from New England BioLabs (U.S.A.). Diethylpyrocarbonate (DEPC)-treated water, S1 nuclease and polyacrylamide containing a ratio of Acrylamide/Bis (19:1, w/w) were purchased from Thermo (U.S.A.). Triethylammonium acetate, hexafluoro-2-propanol and fluorouracil (5-FU) were purchased from Sigma (U.S.A.). Deionized water was prepared using a Millipore Milli-Q Plus system (Millipore, U.S.A.). All reagents used were of analytical grade.

### Affinity purification of tRNAs from NPECS

Affinity purification was performed according to Tsurui *et al*. (45) with minor modification. Briefly, biotin labeled single-strand DNA (antisense) oligonucleotides complementary to tRNA-Val(UAC) (5’-GCCGACCCCCTCCTTGTAAGGGAGGTGCTC-3’) or tRNA-Leu(CAG) (5’-ACGTCCGTAAGGACACTAACACCTGAAGCT-3’) were mixed with total tRNA from *E. coli*, and then denatured at 95°C for 5 min. After incubation at a temperature 5°C lower than the melting temperature (Tm) of each DNA probe for 1.5 h, streptavidin-magnetic beads (Beaverbio, China) were added and incubated for another 30 min. Subsequently, the biotinylated DNA/tRNA coated beads were separated with a magnet, and washed at 40°C. Finally, the magnetic beads were incubated in RNase-free water at 65°C for 5 min to release the immobilized tRNA with probe and then centrifuged at 10,000 g for 1 min. The supernatants were analyzed immediately by UHPLC.

### S1 nuclease hydrolysis

S1 nuclease was added to purified tRNA samples (8 U enzyme/500 ng tRNA) with reaction buffer (2 μL), the final volume adjusted to 20 μL, and incubated at 25°C for 40 min. The reaction was stopped by the addition of EDTA solution (0.5 M, 0.5 μL). The supernatant was collected by centrifugation at 10,000 g for 1 min for UHPLC-MS analysis.

### Urea-polyacrylamide gel electrophoresis (urea-PAGE)

All samples were separated by vertical slab gel electrophoresis (Mini-Protean Tetra System (Bio-Rad, U.S.A.) using 15% urea-polyacrylamide gel. Samples were electrophoresed at 250 V for 1 h at room temperature (25°C), stained with 1 X SYBR Gold nucleic acid gel stain (Thermo) in Milli Q-Plus water for 30 min, followed by imaging using a Bio-Rad imaging system under UV light.

### UHPLC-MS

UHPLC (Agilent 1290 system Agilent, U.S.A.) using an C_18_ column (Acquity UPLC OST, 2.1×100 mm, 1.7 μm i.d., Waters, U.S.A.) at 60°C with a diode array detector. UHPLC-MS was performed using an Agilent 1290 system (Agilent Technologies, U.S.A.), equipped with a vacuum degasser, a quaternary pump, an autosampler, a diode array detector and an Agilent ultrahigh definition 6545 Q-TOF mass spectrometer. Separation was carried out on an ACQUITY UPLC OST C_18_ Column (2.1×100 mm, 1.7 μm i.d., Waters, U.S.A.) at 60°C. tRNAs were separated by eluding the column at a flow rate of 0.2 mL/min with a mobile phase of 100 mM HFIP+15 mM TEAA containing MeOH of the following concentration: 1% (v:v) for 1.5 min, then 1–14% over 1.5-8.3 min, and finally 14–17 % over 8.3-16.5 min. ESI conditions were as follows: gas temperature 320°C, spray voltage 3.5 kV, sheath gas flow and temperature were set as 12 L/min and 350°C, respectively. Fractions corresponding to each chromatographic peak were collected and freeze-dried. For MS experiment, samples were analyzed in negative mode over an *m/z* range of 500 to 3200.

### Cell culture

All cell lines were purchased from American Type Culture Collection (ATCC). 5-FU-resistant LoVo colorectal adenocarcinoma cell line, HCT-8 human ileocecal colorectal adenocarcinoma cell line and its 5-FU-resistant was cultured in RPMI 1640 medium (Gibco, New Zealand) containing 10% fetal bovine serum (FBS), 1% penicillin/streptomycin (P/S) in a humidified 5% CO_2_ atmosphere at 37°C. LoVo colorectal adenocarcinoma cell line was cultured in F-12K medium (Thermo) containing 10% fetal bovine serum (FBS), 1% penicillin/streptomycin (P/S) in a humidified 5% CO_2_ atmosphere at 37°C. All tested RNA samples were dissolved in nuclease free water and stored in −80°C before use. 5-FU was dissolved in dimethyl sulfoxide (DMSO) and used as a positive control.

### Cytotoxicity determination

Cells (5×10^3^ in 100 μL cultured medium) were seeded onto 96-well plates. After 20 h, cells were treated with varying concentrations of RNA sample solutions with Lipofectamine RNAiMAX Transfection Reagent in Opti-MEM medium (Thermo) according to manufacturer’s instructions. Cells without any treatment were used as control, cells treated with liposomes were used treatment control. Cell viability was determined, after 48 h. MTT [3-(4,5-dimethylthiazol-2-yl)-2,5-diphenyltetrazolium bromide] solution (50 μL per well, 1 mg/mL, Thermo) was added to each well and incubated for 4 h at 37°C, followed by DMSO (200 μL) and the A_570_ was measured using a SpectraMax 190 microplate reader (Molecular Devices, U.S.A). Dose-response curves were constructed and the IC_50_ values were calculated by GraphPad Prism 5.0 (GraphPad, U.S.A.). Each experiment was carried out for three times. IC_50_ results were expressed as means ± standard deviation.

### Clonogenic assay

The clonogenic assay was performed according to Franken *et al* (46) with minor modification. Briefly, cells were plated at a density of 1000 cells/well with culture medium in 6-well plates. After 20 h, the medium was changed to medium containing RNA samples (50 nM for HCT-8 and 25 nM for LoVo cells) with liposomal transfection, 5-FU at 50 μM or blank Opti-MEM for a further 48 h. Cells were maintained in normal culture medium for the following 14 days. After fixation for 20 min with a 4% paraformaldehyde fix solution (Beyotime, China), the cells were stained with crystal violet (Beyotime, China) for 10 min. Finally, the number of colonies with more than 50 individual cells were counted using ImageJ software.

### Wound healing assay

Cells (5×10^5^ in 100 μL cultured medium) were grown in 6-well plates and for 20 h until confluent. A scratch was made by using a sterile 1 mL pipette tip and the medium was changed to medium containing RNA samples (50 nM for HCT-8 and 25 nM for LoVo cells) with liposomal transfection or 5-FU at 50 μM. The cells were viewed at 10X objective and photographed using a phase contrast microscope (Leica microsystems, Germany) at various time points (0, 24 and 48 h). ImageJ software was applied to quantify the area of wound created. Wound healing rate was calculated using the formula Wound healing rate = [(Wound area at 0 h – Wound area at 24 or 48 h)/Wound area at 0 h] × 100.

### Simulations of 3D structures of EC83 mimic and its modified derivatives

The initial standard A-form double helix structures for the EC83 mimic and its modified derivatives were built by NAB (47) in Amber18 package (48). The force field used was OL3 (49) (leaprc.RNA.OL3 in Amber18) and the parameters for the modified nucleotides (s^4^U and Gm) were from Aduri *et al* (50). Each system was solvated in a 12Å TIP3P cubic water box (∼6000 water molecules) and then 42 Na^+^ were randomly added to neutralize the whole system.

Before simulations, these systems were minimized in two stages. At the first stagethe RNA molecule was restricted by 20 kcal/mol Å harmonic potential with the energy of the solvent minimized for 8000 steps, where the first 4000 were steepest descent steps and the last 4000 were conjugated gradient steps. At the second stage, another 8000 minimization steps were performed in the same way except that the constraints exerted on the solvate were removed.

After energy optimization, the system was heated from 0 to 300 K and the temperature equilibration was allowed at 300 K for 200 ps. Subsequently, another 200 ps equilibration simulations were performed. The solvate was restricted as described above. Besides, the time step was set to 1 fs and run at NPT ensemble using the parallel version of pmemd (51), with the primary molecular dynamics engine within Amber. 300 ns production simulations was performed for each dsRNA. The temperature was equilibrated at 300 K by Langevin dynamics in the first 100 ns and then gradually increased to 500 K in the last 200 ns with 10 K intervals. The time step was increased from 1 to 2 fs due to the application of SHAKE algorithm. The nonbond cutoff was set to 10 Å and the long-range electrostatic interactions were evaluated by PME method. The pressure was kept around 1 bar (0.987 atm) and all simulations were performed on Nvidia GPU cards by the GPU version of pmemd.

### Comparison of stability of EC83 mimic and its modified derivatives

The stability of these dsRNAs was compared in terms of backbone (P, O3’, O5’, C3’, C4’, C5’) RMSD referred to the stable structures and the extent of hydrogen bondings in 22 base pairs. As the fluctuation of RMSD for each dsRNA was small, the stable structures for these dsRNAs were the average structures in 50-100 ns. Furthermore, the two criterions were calculated by cpptraj (52) in Amber and following data analysis were based on numpy and matplotlib, two popular python packages. The 3D structures were visualized by pymol (53).

### Statistical and data analysis

All experiments results were expressed as mean ± SD. Statistical significance was analyzed using a two-tailed Student t test (GraphPad Prism) or one-way ANOVA followed by post hoc analysis.

## Acknowledgements

This work was financially funded by The Science and Technology Development Fund, Macau SAR (File no. 0023/2019/AKP, 015/2017/AFJ).

## REFERENCES

1. Ley RE, Peterson DA, Gordon JI. 2006. Ecological and evolutionary forces shaping microbial diversity in the human intestine. Cell 124:837–48.

2. Neish AS. 2009. Microbes in gastrointestinal health and disease. Gastroenterology 136:65–80.

3. Marchesi JR, Adams DH, Fava F, Hermes GD, Hirschfield GM, Hold G, Quraishi MN, Kinross J, Smidt H, Tuohy KM, Thomas LV, Zoetendal EG, Hart A. 2016. The gut microbiota and host health: a new clinical frontier. Gut 65:330–9.

4. Gordon DM, Cowling A. 2003. The distribution and genetic structure of Escherichia coli in Australian vertebrates: host and geographic effects. Microbiology 149:3575–86.

5. Bentley R, Meganathan R. 1982. Biosynthesis of vitamin K (menaquinone) in bacteria. Microbiol Rev 46:241–80.

6. Kosek M, Bern C, Guerrant RL. 2003. The global burden of diarrhoeal disease, as estimated from studies published between 1992 and 2000. Bull World Health Organ 81:197–204.

7. Arthur JC, Gharaibeh RZ, Muhlbauer M, Perez-Chanona E, Uronis JM, McCafferty J, Fodor AA, Jobin C. 2014. Microbial genomic analysis reveals the essential role of inflammation in bacteria-induced colorectal cancer. Nat Commun 5:4724.

8. Dejea CM, Fathi P, Craig JM, Boleij A, Taddese R, Geis AL, Wu X, DeStefano Shields CE, Hechenbleikner EM, Huso DL, Anders RA, Giardiello FM, Wick EC, Wang H, Wu S, Pardoll DM, Housseau F, Sears CL. 2018. Patients with familial adenomatous polyposis harbor colonic biofilms containing tumorigenic bacteria. Science 359:592–597.

9. Nougayrede JP, Homburg S, Taieb F, Boury M, Brzuszkiewicz E, Gottschalk G, Buchrieser C, Hacker J, Dobrindt U, Oswald E. 2006. Escherichia coli induces DNA double-strand breaks in eukaryotic cells. Science 313:848–51.

10. Ferlay J, Steliarova-Foucher E, Lortet-Tieulent J, Rosso S, Coebergh JW, Comber H, Forman D, Bray F. 2013. Cancer incidence and mortality patterns in Europe: estimates for 40 countries in 2012. Eur J Cancer 49:1374–403.

11. Siegel R, Desantis C, Jemal A. 2014. Colorectal cancer statistics. CA Cancer J Clin 64:104–17.

12. Cao KY, Pan Y, Yan TM, Jiang ZH. 2020. Purification, characterization and cytotoxic activities of individual tRNAs from Escherichia coli. Int J Biol Macromol 142:355–365.

13. Storz G. 2002. An expanding universe of noncoding RNAs. Science 296:1260–3.

14. Barciszewska MZ, Perrigue PM, Barciszewski J. 2016. tRNA--the golden standard in molecular biology. Mol Biosyst 12:12–7.

15. Fu H, Feng J, Liu Q, Sun F, Tie Y, Zhu J, Xing R, Sun Z, Zheng X. 2009. Stress induces tRNA cleavage by angiogenin in mammalian cells. FEBS Lett 583:437–42.

16. Yamasaki S, Ivanov P, Hu GF, Anderson P. 2009. Angiogenin cleaves tRNA and promotes stress-induced translational repression. J Cell Biol 185:35–42.

17. Thompson DM, Parker R. 2009. The RNase Rny1p cleaves tRNAs and promotes cell death during oxidative stress in Saccharomyces cerevisiae. J Cell Biol 185:43–50.

18. Gebetsberger J, Polacek N. 2013. Slicing tRNAs to boost functional ncRNA diversity. RNA Biol 10:1798–806.

19. Shen Y, Yu X, Zhu L, Li T, Yan Z, Guo J. 2018. Transfer RNA-derived fragments and tRNA halves: biogenesis, biological functions and their roles in diseases. J Mol Med (Berl) 96:1167–1176.

20. Keam SP, Hutvagner G. 2015. tRNA-Derived Fragments (tRFs): Emerging New Roles for an Ancient RNA in the Regulation of Gene Expression. Life (Basel) 5:1638–51.

21. Goodarzi H, Liu X, Nguyen HB, Zhang S, Fish L, Tavazole SF. 2015. Endogenous tRNA-Derived Fragments Suppress Breast Cancer Progression via YBX1 Displacement. Cell 161:790–802.

22. Mo D, Jiang P, Yang Y, Mao X, Tan X, Tang X, Wei D, Li B, Wang X, Tang L, Yan F. 2019. A tRNA fragment, 5′-tiRNA^č^, suppresses the Wnt/ β-catenin signaling pathway by targeting *FZD3* in breast cancer. Cancer Letters 457:60–73.

23. Martinez G, Choudury SG, Slotkin RK. 2017. tRNA-derived small RNAs target transposable element transcripts. Nucleic Acids Res 45:5142–5152.

24. Diebel KW, Zhou K, Clarke AB, Bemis LT. 2016. Beyond the Ribosome: Extra-translational Functions of tRNA Fragments. Biomark Insights 11:1–8.

25. Boccaletto P, Machnicka MA, Purta E, Piatkowski P, Baginski B, Wirecki TK, de Crecy-Lagard V, Ross R, Limbach PA, Kotter A, Helm M, Bujnicki JM. 2018. MODOMICS: a database of RNA modification pathways. 2017 update. Nucleic Acids Res 46:D303–D307.

26. Vainio H, Miller AB. 2003. Primary and secondary prevention in colorectal cancer. Acta Oncol 42:809–15.

27. Szarynska M, Olejniczak A, Kobiela J, Spychalski P, Kmiec Z. 2017. Therapeutic strategies against cancer stem cells in human colorectal cancer (Review). Oncol Lett 14:7653–7668.

28. Dorsett Y, Tuschl T. 2004. siRNAs: applications in functional genomics and potential as therapeutics. Nat Rev Drug Discov 3:318–329.

29. Song J, Yi C. 2017. Chemical Modifications to RNA: A New Layer of Gene Expression Regulation. ACS Chem Biol 12:316–325.

30. Nachtergaele S, He C. 2018. Chemical Modifications in the Life of an mRNA Transcript. Annu Rev Genet 52:349–372.

31. Roundtree IA, Evans ME, Pan T, He C. 2017. Dynamic RNA Modifications in Gene Expression Regulation. Cell 169:1187–1200.

32. Harcourt EM, Kietrys AM, Kool ET. 2017. Chemical and structural effects of base modifications in messenger RNA. Nature 541:339–346.

33. Durdevic Z, Schaefer M. 2013. tRNA modifications: necessary for correct tRNA-derived fragments during the recovery from stress? Bioessays 35:323–7.

34. Phizicky EM, Alfonzo JD. 2010. Do all modifications benefit all tRNAs? FEBS Letters 584:265–271.

35. Corey DR. 2007. Chemical modification: the key to clinical application of RNA interference? J Clin Invest 117:3615–22.

36. Blanco S, Dietmann S, Flores JV, Hussain S, Kutter C, Humphreys P, Lukk M, Lombard P, Treps L, Popis M, Kellner S, Holter SM, Garrett L, Wurst W, Becker L, Klopstock T, Fuchs H, Gailus-Durner V, Hrabe de Angelis M, Karadottir RT, Helm M, Ule J, Gleeson JG, Odom DT, Frye M. 2014. Aberrant methylation of tRNAs links cellular stress to neuro-developmental disorders. EMBO J 33:2020–39.

37. Eckburg PB, Bik EM, Bernstein CN, Purdom E, Dethlefsen L, Sargent M, Gill SR, Nelson KE, Relman DA. 2005. Diversity of the human intestinal microbial flora. Science 308:1635–8.

38. Gill SR, Pop M, Deboy RT, Eckburg PB, Turnbaugh PJ, Samuel BS, Gordon JI, Relman DA, Fraser-Liggett CM, Nelson KE. 2006. Metagenomic analysis of the human distal gut microbiome. Science 312:1355–9.

39. Lax AJ. 2005. Opinion: Bacterial toxins and cancer--a case to answer? Nat Rev Microbiol 3:343–9.

40. Lara-Tejero M, Galán JE. 2000. A Bacterial Toxin That Controls Cell Cycle Progression as a Deoxyribonuclease I-Like Protein. Science 290:354–357.

41. Wang X, Huycke MM. 2007. Extracellular superoxide production by Enterococcus faecalis promotes chromosomal instability in mammalian cells. Gastroenterology 132:551–61.

42. Markowitz VM, Chen IM, Palaniappan K, Chu K, Szeto E, Grechkin Y, Ratner A, Jacob B, Huang J, Williams P, Huntemann M, Anderson I, Mavromatis K, Ivanova NN, Kyrpides NC. 2012. IMG: the Integrated Microbial Genomes database and comparative analysis system. Nucleic Acids Res 40:D115–22.

43. Levitz R, Chapman D, Amitsur M, Green R, Snyder L, Kaufmann G. 1990. The optional E. coli prr locus encodes a latent form of phage T4-induced anticodon nuclease. EMBO J 9:1383–9.

44. Haiser HJ, Karginov FV, Hannon GJ, Elliot MA. 2008. Developmentally regulated cleavage of tRNAs in the bacterium Streptomyces coelicolor. Nucleic Acids Res 36:732–41.

45. Tsurui H, Kumazawa Y, Sanokawa R, Watanabe Y, Kuroda T, Wada A, Watanabe K, Shirai T. 1994. Batchwise purification of specific tRNAs by a solid-phase DNA probe. Anal Biochem 221:166–72.

46. Franken NA, Rodermond HM, Stap J, Haveman J, van Bree C. 2006. Clonogenic assay of cells in vitro. Nat Protoc 1:2315–9.

47. Macke TJ, Case DA. 1998. Modeling unusual nucleic acid structures. Molecular Modeling of Nucleic Acids 682:379–93.

48. Case DA, Cheatham TE, Darden T, Gohike H, Luo R, Merz KM, Onufriev A, Simmerling C, Wang B, Woods RJ. 2005. The Amber biomolecular simulation programs. J Comput Chem 26:1668–88.

49. Zgarbová M, Otyepka M, Sponer J, Mládek A, Banáš P, Cheatham TE, Jurečka P. 2011. Refinement of the Cornell et al. nucleic acids force field based on reference quantum chemical calculations of glycosidic torsion profiles. J Chem Theory Comput 7:2886–2902.

50. Aduri R, Psciuk BT, Saro P, Taniga H, Schlegel HB, SantaLucia J. 2007. Amber force field parameters for the naturally occurring modified nucleosides in RNA. J Chem Theory Comput 3:1464–75.

51. Salomon-Ferrer R, Gotz AW, Poole D, Grand SL, Walker RC. 2013. Routine microsecond molecular dynamics simulations with Amber on Gpus. 2. Explicit solvent particle Mesh Ewald. J Chem Theory Comput 9:3878–88.

52. Roe DR, Cheatham TE. 2013. Ptraj and Cpptraj: software for processing and analysis of molecular dynamics trajectory data. J Chem Theory Comput 9:3084–95.

53. DeLano WL. 2002. Pymol: an open-source molecular graphics tool. CCP4 Newsletter on protein crystallography 40:82–92.

